# The high-quality genome assembly of Coelioxoides waltheriae (Apidae: Nomadinae) reveals gene family dynamics and evolutionary shifts related to its cleptoparasitic lifestyle

**DOI:** 10.64898/2026.05.18.725936

**Authors:** Felipe Cordeiro Dias, Heraldo Mauch, Paulo Cseri Ricardo, Beatriz Tanzi Martins, Natalia de Souza Araujo, Maria Cristina Arias

## Abstract

Cleptoparasitism, or brood parasitism, is a striking behavioral strategy observed in approximately 13% of all bee species, yet its genomic underpinnings remain largely unexplored. We present the first high-quality genome assembly of the Neotropical cleptoparasitic bee *Coelioxoides waltheriae* (Nomadinae), a species that parasitizes the nests of *Tetrapedia diversipes*. The final assembly comprises 194.8 Mbp across 388 contigs, with an N50 of 1.47 Mbp and 97.4% BUSCO completeness, representing the second smallest genome among cleptoparasitic bees. Repetitive elements constitute only 14.6% of the genome, suggesting that its compact size is primarily driven by repeat reduction rather than gene loss. Comparative genomic analyses across 42 hymenopteran species revealed a pronounced contraction bias in gene family size changes in *C. waltheriae* (expansion ratio of 13.66%), a pattern also observed in other cleptoparasitic lineages. Expanded orthogroups were enriched for cuticle-related genes (e.g., *PiggyBac* transposases) potentially linked to host infiltration and defense, while contracted orthogroups showed significant reductions in sensory perception (e.g., odorant receptors), detoxification (e.g., cytochrome P450), and metabolic genes, reflecting the reduced ecological demands of a parasitic lifestyle. Furthermore, non-target DNA analysis identified associations with *Roubikia* mites (a known symbiont of its host), as well as fungi and bacteria, providing ecological context for this species. Our findings establish a critical genomic reference for cleptoparasitic bees, demonstrating that the evolution of parasitism is associated with targeted gene family contractions in sensory and metabolic functions alongside expansions in cuticle and transposable element-related genes, offering new insights into the genomic signatures of behavioral specialization.

## Introduction

Bees are the most important group of pollinators worldwide, largely due to their foraging activities, which involve the collection of floral resources such as pollen, nectar, and oils (Michener 2007, Jogesh et al. 2016, Khalifa et al. 2021). Throughout their evolutionary history, bees have diversified not only in morphology and ecology but also in behavior, giving rise to a wide range of reproductive and foraging strategies. From an evolutionary perspective, the transition from predatory ancestors to a pollen/nectar-based diet represents a major ecological shift that shaped the diversification of bees. This change, tightly associated with the rise and radiation of angiosperms during the mid-Cretaceous, promoted not only morphological and physiological innovations, but also a remarkable diversification of life-history strategies and behaviors (Ollerton 2017; Murray et al. 2018). While pollen collection and nest provisioning remain the dominant conditions among extant bees, convergent departures from this ancestral foraging strategy highlight the evolutionary lability of bee behavior and the potential for alternative reproductive strategies to arise under specific ecological and selective pressures (Ollerton 2017; Dorey and Schiestl 2024). Among these, parasitic behaviors represent some of the most striking deviations from the typical pollen-collecting lifestyle (Goulson and Hughes 2015).

In ecological terms, parasitism involves interactions in which one species benefits at the expense of another (Phillips 2012). Among bees, cleptoparasitism, also referred to as brood parasitism, is the most remarkable manifestation of this strategy. In this behavior, female parasites invade host nests to lay their eggs, and the emerging larvae consume the stored provisions, ensuring their development. Thus, the parasite completes its life cycle without investing in nest construction or resource collection for the offspring (Michener, 2007).

Approximately 13% of all described bee species are obligate cleptoparasites, distributed across four of the seven bee families, suggesting multiple independent evolutionary origins of this strategy (Danforth et al. 2019; Ricardo et al. 2024). By eliminating the need for nest construction and active foraging, cleptoparasitism may reduce energetic and ecological constraints associated with reproduction (Michener, 2007). The high frequency of cleptoparasitism in bees has been linked to the early evolution of nest provisioning behavior, particularly the storage of pollen and other floral resources within brood cells (Sheffield et al. 2013). This ecological context likely created repeated opportunities for the exploitation of host nests, facilitating the recurrent emergence of cleptoparasitic strategies across distantly related lineages. As a result, cleptoparasitism in bees represents one of the most striking examples of convergent behavioral evolution within a single animal clade. Beyond individual advantages, cleptoparasitic bees may also play complex roles in natural communities by influencing host population dynamics and modulating competition among solitary bees (Sheffield et al. 2013). While the behavioral and ecological aspects of cleptoparasitic bees have received scientific attention, their genomic underpinnings remain largely unexplored. As of December 2025, only a small fraction of the 181 bee reference genomes available at NCBI correspond to cleptoparasitic species, including representatives of the genera *Coelioxys, Epeolus, Holcopasites, Sphecodes*, and *Stelis* ((Sayers et al. 2024), along with additional *Nomada* species generated by the Darwin Tree of Life project (Darwin Tree of Life Project Consortium, 2022). This data limitation has been an constraint on addressing comparative analyses on the genetic signatures associated with parasitic behavior and its evolution.

Comparative genomic studies in parasitic Hymenoptera, including parasitoid wasps (Ye et al. 2024) and parasitic ants (Borowiec et al. 2021), have revealed some genomic patterns in parasites, such as changes in the genomes sizes, often driven by variation in repetitive DNA content, although some parasitic species does not seen to have their genome size influenced by the proportion of repetitive elements (Sless, Searle, et al. 2022). These patterns suggest that shifts toward parasitic lifestyles may be accompanied by genome changes across independent lineages. In bees, a dynamic of genome size change linked to repetitive elements have been proposed, as exemplified by orchid bees (Apidae: Euglossini), which possess comparatively large genomes associated with expanded repetitive elements (Brand and Ramírez 2017, Walsh et al. 2022). Together, these observations highlight repetitive element dynamics as a relevant component of genome evolution in Hymenoptera, though their role may not be restricted to genome size changes alone.

*Coelioxoides waltheriae* Ducke, 1908 is a cleptoparasitic bee, native to the Neotropical region and represents a compelling model for studying the genomic basis of parasitic lifestyles. Its primary host, the solitary bee *Tetrapedia diversipes* Klug, 1810, commonly nests in pre-existing cavities, including trap-nests, facilitating direct observation of parasitism events (Alves-dos-Santos et al. 2002). Although *Coelioxoides* and *Tetrapedia* were historically considered close relatives within the tribe Tetrapediini, recent phylogenomic analyses have repositioned *Coelioxoides* within the predominantly cleptoparasitic clade Nomadinae (Sless, Branstetter, et al. 2022). This revised phylogenetic placement highlights cleptoparasitism as a deeply conserved and evolutionarily significant trait within the Nomadinae lineage.

In this context, genomic data represent a critical resource for investigating the evolutionary association between cleptoparasitism and genome evolution in bees. Building on the first omic investigation of a *Coelioxoides* species (Ricardo et al. 2024), we present the first high-quality genome assembly and comprehensive annotation of *Coelioxoides waltheriae*, establishing a new genomic reference for cleptoparasitic bees. Beyond genome structure and composition, we integrate analyses of repetitive elements, orthology inference, phylogenomics, and gene family expansions and contractions across Anthophila to investigate the evolutionary trajectories associated with parasitism. In addition, we leverage non-target DNA recovered from long-read sequencing of *C. waltheriae* to characterize associated microorganisms, fungi, and plants, providing an ecological and evolutionary context for its biology. Together, these approaches provide a genomic framework to disentangle the evolutionary interplay between cleptoparasitism, genome architecture, and ecological interactions in bees.

## Results

### Genome Assembly and Structure

The PacBio HiFi sequencing resulted in 601,916 reads, totaling 6,786,696,209 bases (Table 1). GenomeScope analysis estimated a genome size of ∼196 Mbp, predominantly composed of unique sequences (88.9%), with low heterozygosity (0.818%), a low error rate (0.434%), and an average coverage of 21x (Supplementary Fig. S1). The initial assembly generated with Flye comprised 196,294,652 bp, with an N50 of 1,436,209 and 756 contigs. After haplotype purging, scaffolding, and polishing, the final assembly comprised 194,795,700 bp in 388 contigs, with an N50 of 1,468,442 (Table 1). After the check for mean coverage, all contigs were kept, since all of them presented mean coverage >1. The BlobToolKit analyses (Fig. 1 ; Supplementary Fig. S2) did not detect any contigs associated with contamination. When assessed with the hymenoptera_odb10 dataset (5,991 BUSCO orthologs), the *C. waltheriae* genome displayed 97.4% completeness (97.2% as single-copy and 0.2% duplicated), along with 0.7% fragmented and 1.8% missing orthologs. These values are consistent with high-quality bee genome assemblies (Table 1).

**Fig 1.**
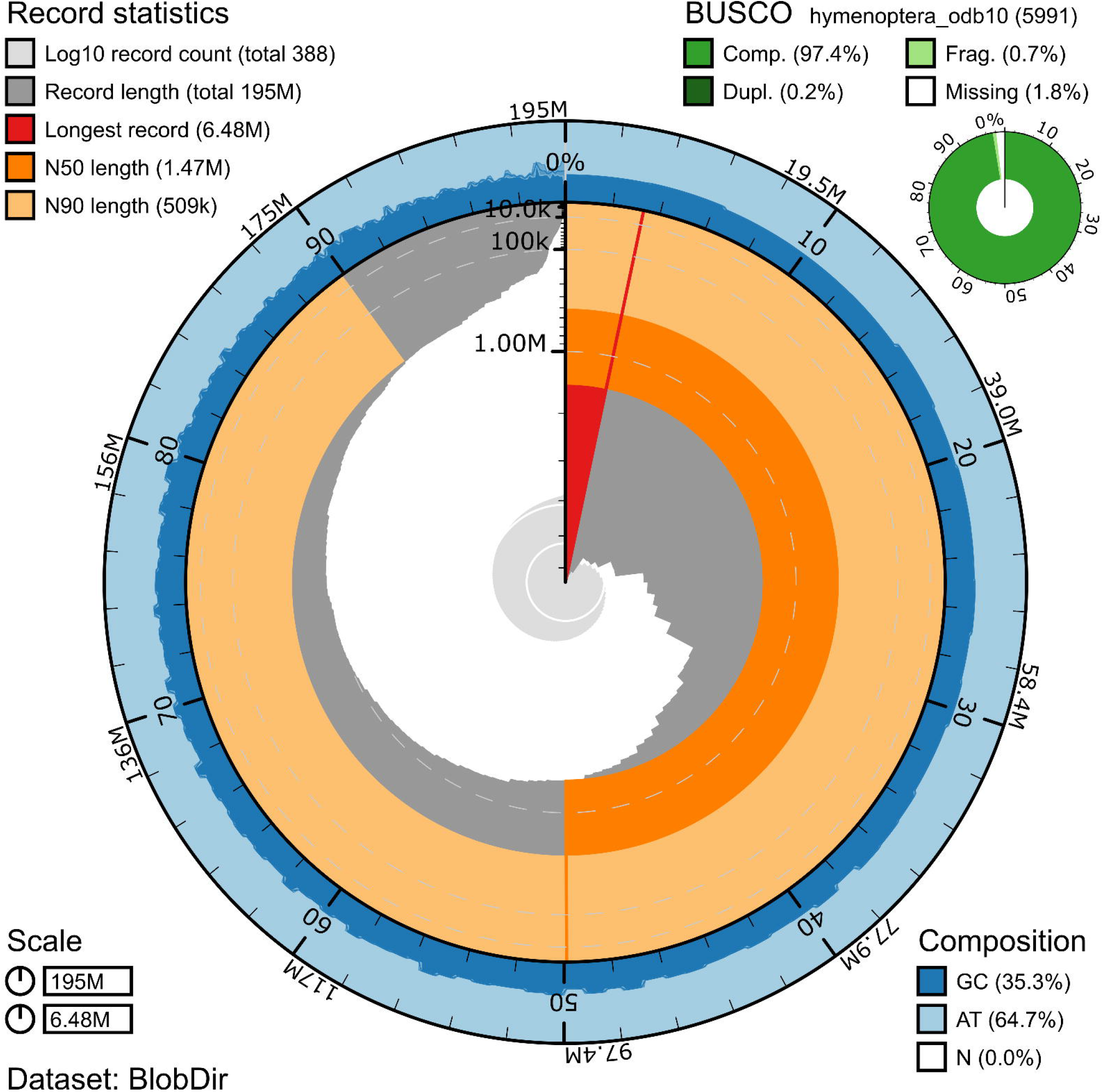

**Table 1.** Overview of Sequencing and Assembly Results for *C. waltheriae*.

| Step/Analysis | Metric | Value |
| --- | --- | --- |
| PacBio HiFi sequencing | Number of reads | 601,916 |
|  | Total bases | 6,786,696,209 |
|  | Estimated coverage | 21x |
| Initial assembly (Flye) | Total size | 196,294,652 bp |
|  | Contigs | 756 |
|  | N50 | 1,436,209 |
| Final assembly | Total size | 194,795,700 bp |
|  | Contigs | 388 |
|  | N50 | 1,468,442 |
|  | L50 | 31 |
|  | L90 | 117 |
|  | Largest scaffold | 6.84 Mbp |
| BlobToolKit | Contamination detection | None |
| BUSCO (hymenoptera_odb10, 5,991 genes) | Complete single-copy | 97.2% |
|  | Complete duplicated | 0.2% |
|  | Fragmented | 0.7% |
|  | Missing | 1.8% |

*C. waltheriae* has the sixth smallest bee genome and the second smallest among cleptoparasitic species. Across the 149 reference bee genomes available in the NCBI (December 2025), together with the *C. waltheriae*, the mean genome size was 323.34 Mbp and the median was 297 Mbp. This similarity between the mean and median suggests a relatively balanced distribution of genome sizes, indicating that extreme values or outliers (e.g., *Xylocopa violacea*, with a genome size higher than 1 Gbp) do not strongly skew the dataset (Supplementary Fig. 2; Supplementary Table 1).

### Repetitive Elements

Repetitive elements comprise 14.6% of the genome, with the majority assigned to the “Unclassified” category (58.29%), followed by Class II elements (20.14%) and Simple Repeats (12.9%). The complete distribution of repetitive element categories is shown in Fig. 2 and Supplementary Table S3. The repetitive elements were presented in approximately 12% of both upstream and downstream gene regions. The majority of genes (95,3%) had at least part of their regulatory regions containing repeats. Additionally, 14.38% and 13.98% of genes had repeats exclusively in the upstream or downstream region, respectively (Supplementary Table S4). Considering its genome size, *C. walteriae* shows a relatively low proportion of repetitive sequences (Fig. 3).

**Fig 2.**
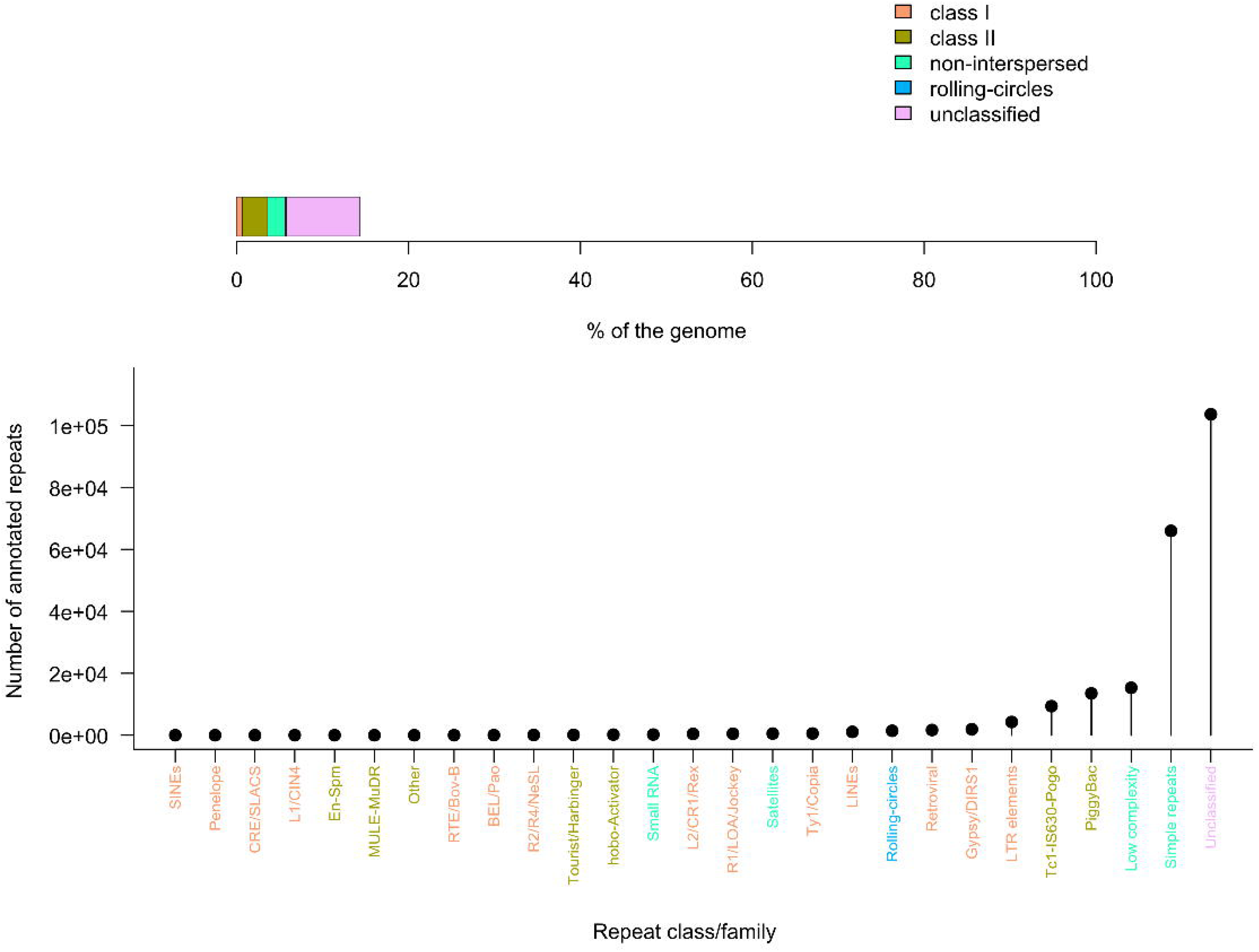

**Fig 3.**
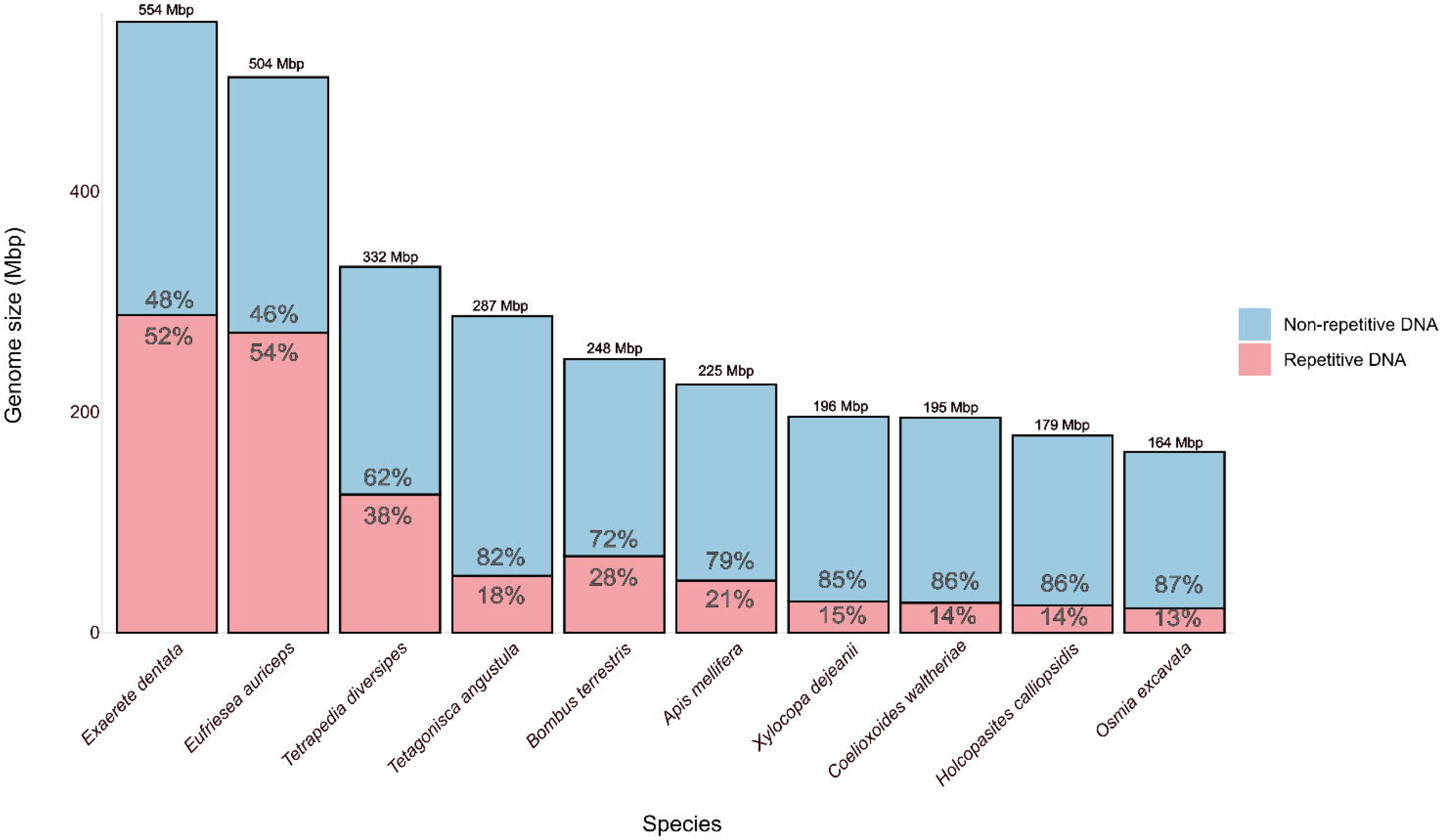

### Genome Annotation

The analysis predicted 11,684 genes and 97 tRNAs, totaling 17,826,909 bp (9.5% of the genome). According to L90 metrics, 87.02% of genes were located within the 117 largest scaffolds. Conversely, 129 scaffolds did not contain any genes (Supplementary Table S6) and corresponded to the shortest scaffolds in the assembly. Functional annotation covered approximately 93% of the predicted genes. Gene Ontology (GO) Biological Process annotations were assigned to 7,199 genes using eggNOG-mapper, yielding a total of 19,781 distinct GO terms (Supplementary Table S5).

### Orthology and genomic comparison

OrthoFinder analysis was run using 467,149 predicted protein sequences from 42 species. Of these, 433,893 sequences (92.9%) were assigned to 17,300 orthogroups, indicating a high degree of gene sharing across taxa, overall high-quality genome annotation and evolutionary common pathways. In addition, 1,029 species-specific orthogroups were identified, comprising 3,428 protein sequences. These lineage-specific groups accounted for just 0.7% of all genes, suggesting a limited number of unique gene innovations per species, as expected in broad orthology assessments involving closely related species (Fig. 4).

**Fig 4.**
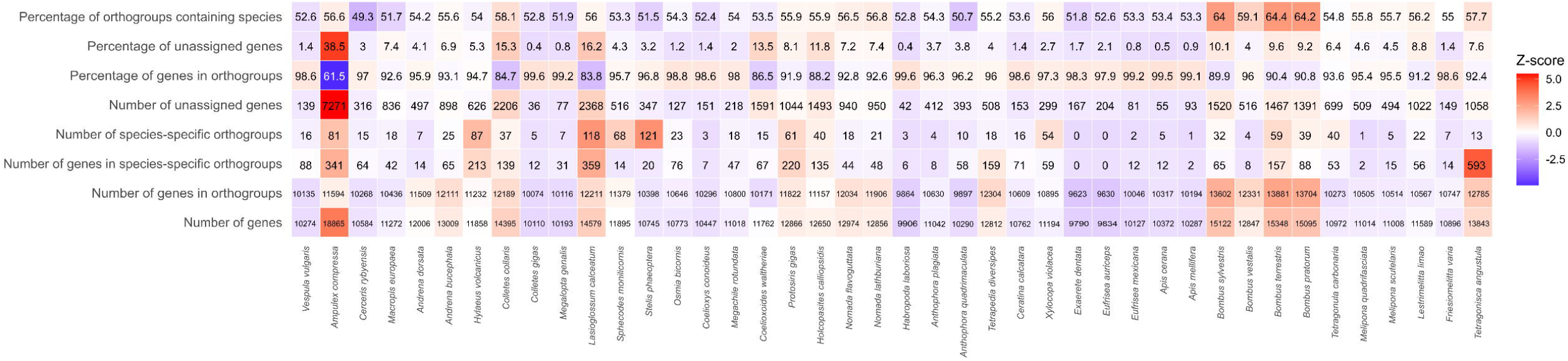

Overall, most species showed a high proportion of proteins clustered in orthogroups, regardless of their total gene count. Several species with compact genomes, such as *Apis mellifera, Megalopta genalis*, and *Melipona scutellaris*, had over 95% of their proteins assigned to orthogroups. In contrast, species with larger gene sets, such as *Ampulex compressa* and *Cerceris rybyensis*, showed lower proportions of assigned proteins (around 60–70%), suggesting the presence of lineage-specific or poorly conserved genes (Fig. 5).

**Fig 5.**
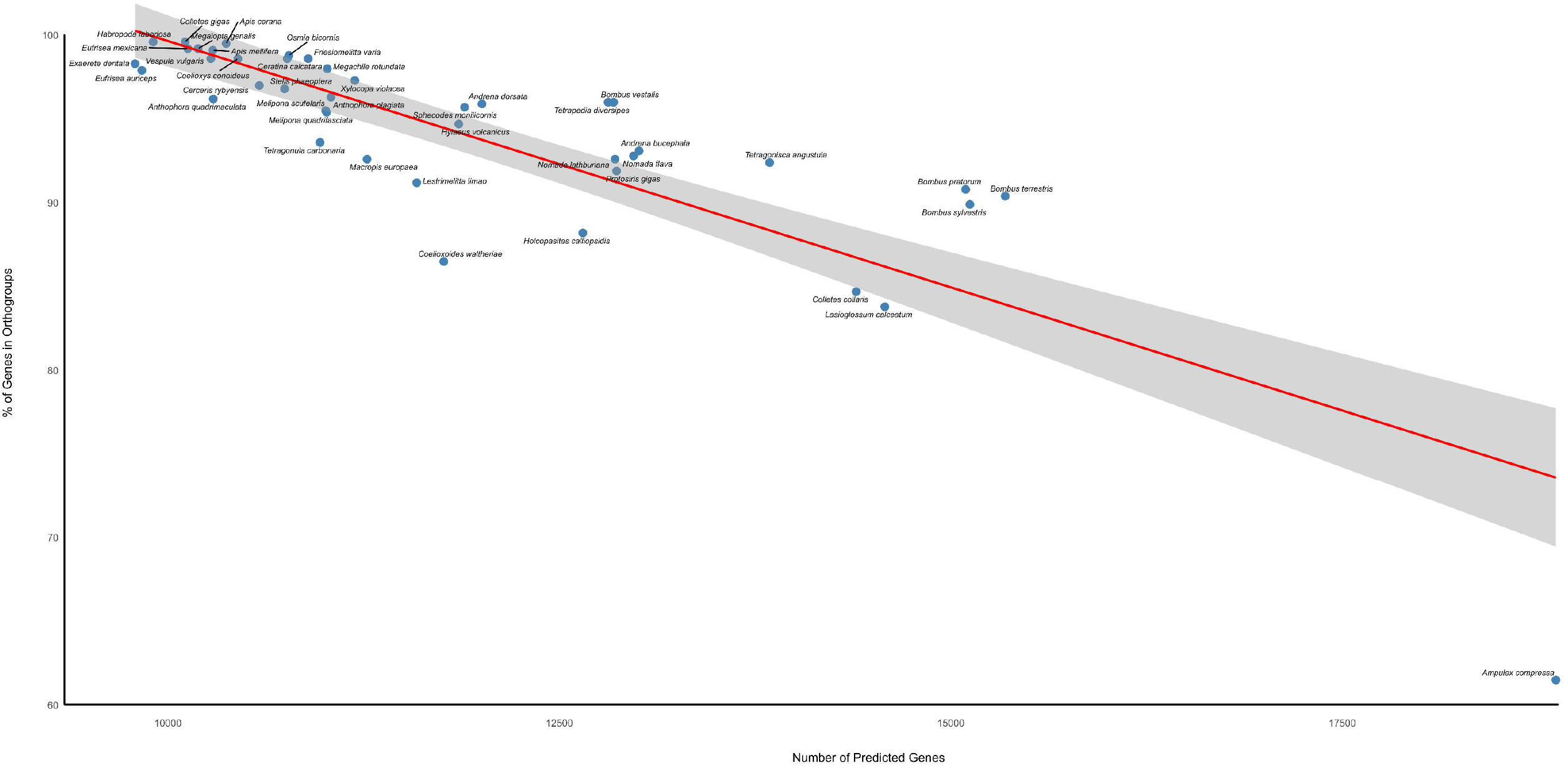

### Gene families size changes

The phylogenetic reconstruction of gene family size evolution across the analyzed species, considering both expansion and contraction events inferred by CAFE5, revealed a slight predominance of contractions throughout the phylogeny. When all branches were considered, contraction events accounted for approximately 54% of the total gene family size changes, whereas expansions represented about 46% (Supplementary Table S7). These patterns were visualized by mapping the expansion ratio for each lineage onto the phylogeny, calculated as the proportion of expansion events relative to the total number of changes (Exp Ratio (%) = (I × 100) / (I + D)), where *I* corresponds to increases and *D* to decreases in gene copy number (Fig. 6).

**Fig 6.**
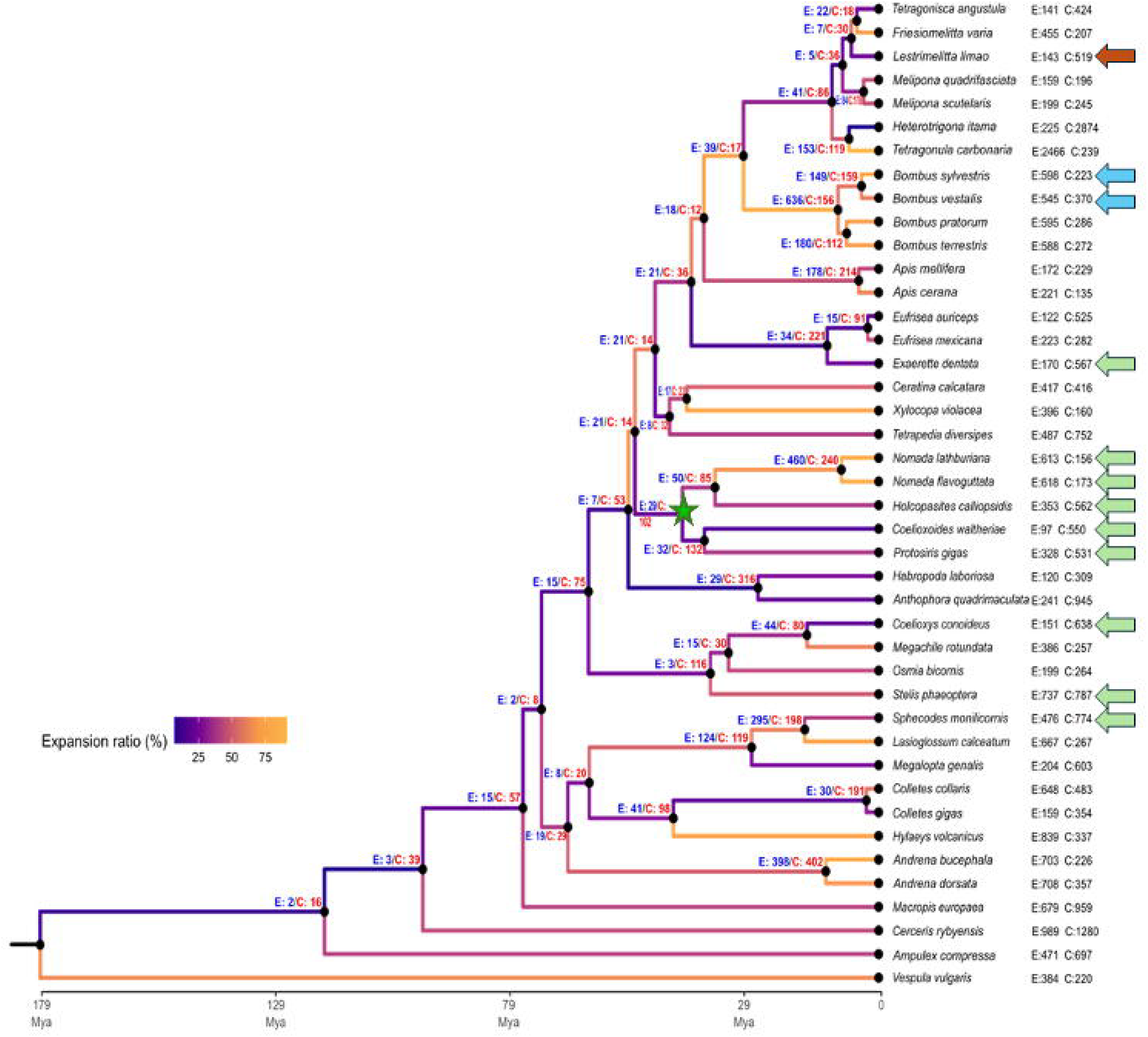

Across most lineages, the relative balance (e.g. the number of contractions and expansions divided by the global number of events per species/node) between expansions and contractions remained close to parity, with moderate variation in expansion ratios among taxa. However, a pronounced shift toward contraction was observed in some cleptoparasitic lineages. In these taxa, decreases in gene copy number substantially exceeded increases, resulting in lower expansion ratios compared to the majority of non-parasitic species included in the analysis (Fig. 6).

The strongest contraction bias among all sampled species was detected in *Coelioxoides waltheriae* (Fig. 6). This lineage exhibited 112 gene family expansions and 568 contractions, corresponding to an expansion ratio of 13.66%, the lowest value observed across the phylogeny. Similar contraction-biased patterns were also present in other cleptoparasitic species, including *Holcopasites calliopsidis* (348 expansions vs. 586 contractions) and *Coelioxys conoideus* (150 expansions vs. 641 contractions), both of which displayed substantially reduced expansion ratios relative to most other taxa. Overall, these results indicate that although gene family expansions and contractions occur throughout the phylogeny, contraction events represent the dominant form of gene family size change in several cleptoparasitic lineages, particularly in *C. waltheriae*, where reductions in gene copy number are markedly more frequent than increases (Fig. 6).

Of the 680 orthogroups (OGs) showing gene copy number changes in *C. waltheriae*, 16.5% corresponded to expansions and 83.5% to contractions. Approximately half of these OGs were functionally annotated based on similarity searches and domain assignments (Supplementary Table S8). Among the OGs showing gene copy changes, 298 OGs were identified as significantly expanded or contracted in the CAFE5 analysis, including 34 expansions (∼30% of all expanded OGs) and 264 contractions (∼46% of all contracted OGs).

Functional enrichment analyses, performed separately for expanded and contracted OGs, revealed a clear asymmetry in the distribution of biological functions between these two sets (Fig. 7). Among the expanded OGs with assigned functions, two relevant categories emerged in the context of cleptoparasitic lifestyles. The first comprised genes associated with structural cuticle components, represented by two orthogroups containing proteins annotated with cuticle-related domains. The second category comprised transposable elements–derived functions, including 2 orthogroups associated with PiggyBac transposases and retrovirus-related pol polyprotein families (Supplementary Table S8) (Fig. 7).

**Fig 7.**
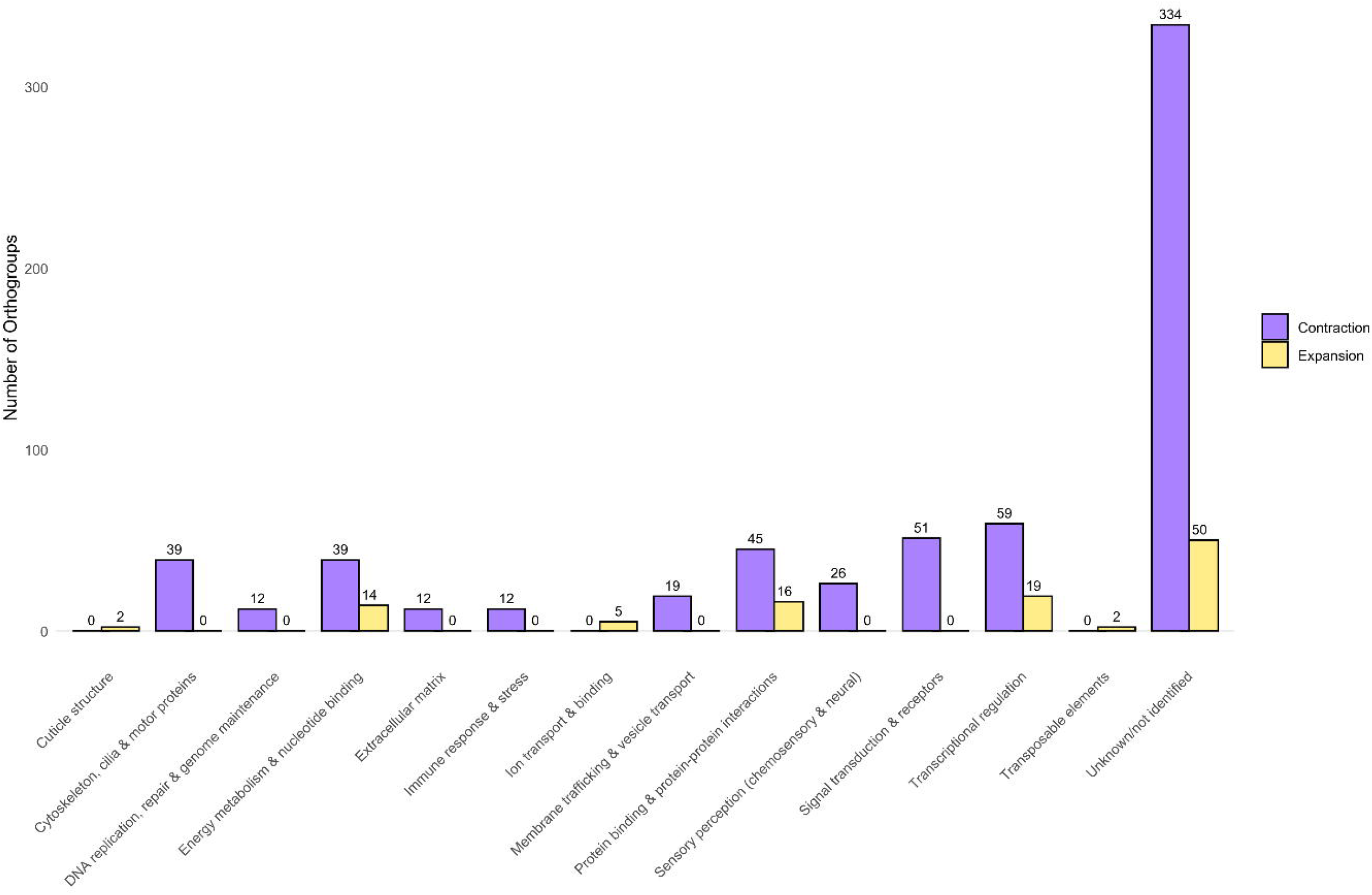

In contrast, contracted OGs with functional annotation were distributed across several biological categories and represented a substantially larger portion of the detected gene copy number changes. A prominent component of these contractions involved genes related to sensory perception. In particular, multiple odorant receptor (OR) orthogroups exhibited reduced copy numbers, consistent with contraction within this receptor family. Contractions were also observed in detoxification-related genes, notably within cytochrome P450 (CYP) orthogroups associated with xenobiotic metabolism. Beyond these functional classes, additional contracted OGs corresponded to genes involved in a broad range of molecular and cellular functions. These included orthogroups annotated with ATP-binding and protein-binding domains, as well as genes associated with ion transport, immune response and stress-related processes (here treated as a single functional category), and enzymes participating in diverse metabolic pathways. Together, these contracted orthogroups encompass proteins involved in general cellular maintenance, transport functions, and multiple aspects of cellular metabolism (Fig. 7).

GO enrichment analyses of contracted orthogroups revealed a strong overrepresentation of functions associated with neural and sensory systems (Fig. 8). The most significantly enriched categories included axon guidance, neuron projection development, neurogenesis, neuron differentiation, and nervous system development, together with processes related to cell morphogenesis and synapse organization. Sensory-related functions were also enriched, particularly odor perception, chemical stimulus detection, and behavioral processes, consistent with the contraction observed in odorant receptor orthogroups. Additionally, several enriched terms were associated with ion transport and neuronal signaling, including voltage-gated potassium channel activity, transporter complexes, and plasma membrane-associated components. Developmental categories such as metamorphosis, wing morphogenesis, and imaginal disc development were likewise recurrently enriched, suggesting that contractions disproportionately affected genes involved in neuronal connectivity, sensory perception, signaling, and developmental regulation. In contrast, GO enrichment analyses of expanded orthogroups identified only four significantly enriched categories, primarily associated with mitotic and chromosomal processes. These included mitotic metaphase chromosome alignment, mitotic chromosome movement toward spindle poles, and positive regulation of spindle microtubule attachment, together with cellular response to iron ions. Overall, the enrichment profile of expanded orthogroups suggests a more restricted functional pattern centered on cell division and chromosome segregation dynamics (Supplementary Table S9).

**Fig 8.**
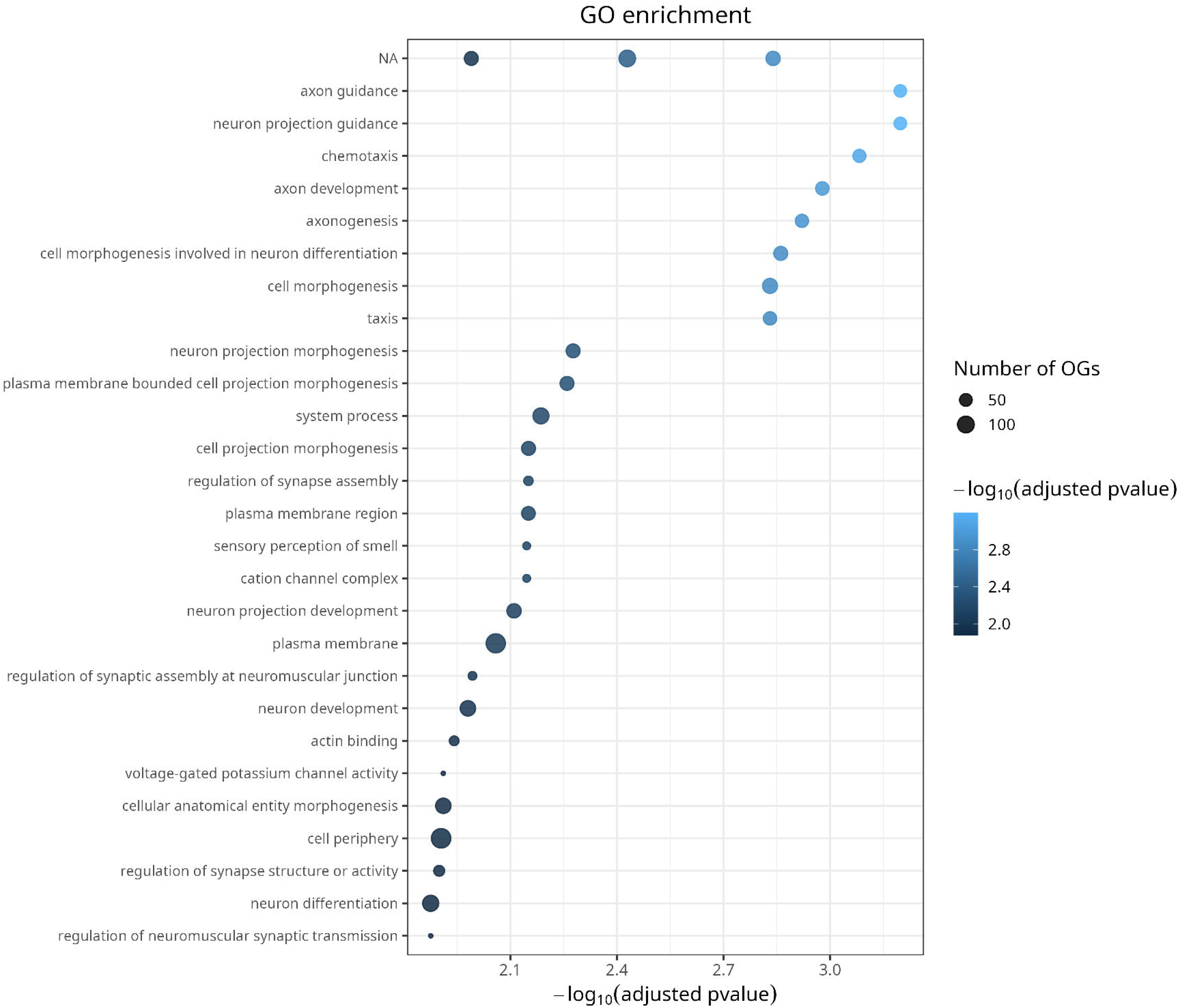

### Non-target DNA sequences identification

Out of the 82,056 reads that did not align to the masked genome, 1,883 aligned against a nucleotide database sequence (NCBI). A total of 157 hits were verified as Acari (14 species). Most of the mite species identified were generalists, such as *Dermatophagoides sp*. (85 reads) and *Sarcoptes scabiei* (41 reads), which are also human-associated (Querner et al., 2024). Notably, two reads were assigned to the genus *Roubikia*, mites associated with the bee genera *Tetrapedia* and *Coelioxoides* (Klimov, et al., 2007; Cordeiro et al., 2011).

Seven sequences matched fungi, with each read assigned to a specific species. Four of them were plant pathogens (*Fulvia fulva, Botrytis cinerea, Lasiodiplodia theobromae*, and A*spergillus nomiae*), while the other three were human-associated (*Cokeromyces recurvatus, Malassezia globos*a, and *Trichosporon inkin*) (Findley & Oh, 2013; Santos et al., 2022).

A total of 176 hits were classified as bacteria, corresponding to 62 species. Eight species (29 reads) were classified as bee host-associated: *Bifidobacterium sp*., *Bombella sp*., *Lactobacillus sp*., *Melissococcus plutonius, Parasaccharibacter apium, Parasaccharibacter sp*., *Snodgrassella alvi*, and *Snodgrassella communis*. Three species (11 reads) corresponded to insect (non-Anthophila) sequences: *Aristophania vespae, Oecophyllibacter saccharovorans*, and *Proteus vulgaris*. Furthermore, 80 sequences matched bacteria isolated from human hosts, 18 from plant-associated hosts, and 38 from environmental bacteria or other sources.

Among the remaining sequences, 188 reads aligned to human sequences, suggesting potential environmental or handling contamination. A total of 1,105 were identified as belonging to Hymenoptera, the majority of which (672 reads) corresponded to the Nomadinae subfamily, the same subfamily as *C. waltheriae*. These reads may represent lower-quality sequences that failed to meet the alignment thresholds against the assembled *C. waltheriae* genome. Finally, 242 reads were classified as unassigned or belonging to diverse taxa. No viral sequences were identified in the dataset.

## Discussion

Our study demonstrates that *C. waltheriae* has a relatively small genome compared to the average among bees (Supplementary Fig. 2). The low proportion of repetitive elements (Fig. 2) likely contributes to this feature, and is consistent with observations in other insects and across diverse animal taxa (Tsutsui et al. 2008; Kapusta et al. 2017; Yuan et al. 2024). This suggests that variation in the proportion of repetitive elements may be an important factor shaping genome size.

Bees play a central role as primary pollinators in natural and agricultural ecosystems, making them a key group for evolutionary and genomic research. However, genomic resources remain limited, with fewer than 1% of described species currently represented by sequenced genomes and a strong concentration of available data in the genera *Apis* and *Bombus*, according to the NCBI database. Genomic information becomes even scarcer when considering species with specialized behavioral strategies such as cleptoparasitism, for which only a few species - mostly within the genus *Nomada* - have genomic data available (Odanaka(Odanaka 2024; Martins et al. 2025). Cleptoparasitic bees, including *Coelioxoides waltheriae*, therefore remain largely understudied from genetic, molecular, and evolutionary perspectives. Nevertheless, recent studies have begun to shed light on the molecular basis of their ecological and behavioral specializations, revealing that differences in gene expression underlie key adaptations in this species (Ricardo et al., 2024), as well as in other parasitic bees (Sless, Searle, et al. 2022; Martins et al. 2025; Ricardo et al. 2025).

The high proportion of unclassified repetitive regions may reflect species-specific patterns that differ from those found in families described in the literature. Across the sampled bee genomes, species with smaller genomes (including *C. waltheriae*) tend to show less than 20% repetitive content, while larger genomes tend to contain around 50% repetitive sequences (Fig. 3). Variation in genome size among bees is largely driven by differences in repetitive DNA content rather than by extensive gene loss (Brand et al., 2017; Kapusta et al., 2017). In *C. waltheriae*, the overall gene content is comparable to that of other bee species (∼12,000 genes; NCBI, 2025), indicating that its reduced genome size is not associated with substantial gene depletion. Instead, genome compaction appears to reflect a lower proportion of repetitive elements. From a phylogenetic perspective, this reduction in genome size may represent a relatively recent event. While species of the genus *Nomada* typically possess genomes exceeding 225 Mbp, another Nomadinae species, *Holcopasites calliopsidis*, also exhibits a reduced genome size (Sless, Searle, et al. 2022). This pattern may indicate that genome contraction occurred after the divergence of the tribe Nomadini from other members of the subfamily. Although repetitive sequences are often associated with regulatory innovation and epigenetic modulation (Shapiro and von Sternberg 2005), most regulatory regions in *C. waltheriae* are composed predominantly of non-repetitive sequences, suggesting that genome reduction does not necessarily entail reduced regulatory complexity.

The annotation and orthology analyses provide insights into how ecological specialization may be linked to the genome profile. The concentration of genes within the largest scaffolds indicates that gene-rich regions are well represented in the most contiguous portions of the assembly. In contrast, smaller scaffolds are predominantly composed of repetitive sequences and lack annotated genes, suggesting that they may correspond to repeat-rich or fragmented genomic regions. This pattern is consistent with known challenges in assembling highly repetitive sequences and likely reflects technical limitations of scaffolding rather than true gene absence. Similar patterns of gene distribution, with a small fraction of scaffolds lacking gene content, have been observed in other bee genomes and are often associated with assembly limitations or high repetitive content (Sadd et al. 2015).

The high proportion of genes assigned to orthogroups across most species indicates strong conservation of core gene sets (Fig. 5), consistent with the notion that genome compaction in *C. waltheriae* predominantly affects non-coding and repetitive regions rather than core gene content, a pattern also observed in other insect taxa with specialized lifestyles (Kapusta et al., 2017; (Kapusta et al. 2017; Sproul et al. 2023; Yuan et al. 2024).

The changes in orthogroup gene numbers for *C. waltheriae* revealed a pronounced contraction bias, the strongest observed among all sampled species, alongside contraction-biased patterns in other cleptoparasitic taxa such as Holcopasites calliopsidis and Coelioxys conoideus, suggesting that gene family reduction may be a recurrent feature of the cleptoparasitic lifestyle across independent lineages. Functional analysis revealed that expanding orthogroups in *C. waltheriae* are enriched for nuclear, membrane, and transcription-related functions, while contracted orthogroups are associated with housekeeping, metabolic, and sensory processes, including ATP binding, protein binding, and ion channels (Fig. 7). This pattern suggests targeted genomic changes supporting regulatory complexity and cellular functions, alongside reductions in metabolic and sensory capacities consistent with cleptoparasitic lifestyle.

The expansion in two orthogroups (OG0000044 and OG0009069 – Supplementary Table 8), both linked to cuticle development, may strongly indicate an adaptation to a cleptoparasitic lifestyle. Compared to non-parasitic species, many cleptoparasites tend to have a thicker, more sclerotized cuticle (Michener 2007; Danforth et al. 2019), which can provide additional protection, either by masking the parasite’s scent or defending against host attacks during egg laying. Ricardo et al. (2024) reported upregulation of the *Hr38* nuclear factor in *C. waltheriae*. Among its regulatory roles, this factor has been associated with cuticle gene expression and maintenance of cuticular integrity in *Drosophila* (Kozlova et al. 2009). Our findings broaden the understanding of the cuticle’s role in species adaptation, particularly in relation to the cleptoparasitic lifestyle of *C. waltheriae*.

The observed expansion of gene families encompassing transposable elements, such as PiggyBac (OG0000002) and the retrovirus-related Pol polyprotein (OG0000042), suggests that molecular domestication of transposable elements may contribute to genomic innovation underlying cleptoparasitism in *C. waltheriae*. Molecular domestication, which inactivates transposition and can convert transposable elements into functional genes through changes in coding or regulatory regions, may represent a mechanism contributing to genomic innovation (Bouallègue et al. 2017). In a parasite that must continuously coevolve to overcome host defenses, retaining such elements or their derivatives could provide adaptive advantages by generating genomic diversity (Volff 2006; Sinzelle et al. 2009). Additionally, our results corroborate the transcriptomic evidence of upregulation of PiggyBac genes (Ricardo et al., 2024) and expansion of Retroviral related genes in *Holcopasites calliopsidis*, another cleptoparasite from the same subfamily (Sless, Searle, et al. 2022).

Gene contractions in *C. waltheriae* were particularly pronounced in families associated with sensory perception, chemical defense, and energy metabolism. Several odorant receptor (OR) orthogroups, including OG0004306, OG0000146, OG0009076, OG0000035, OG0009509, and OG0000490, showed copy number reductions, supporting the hypothesis that a parasite can survive having a more limited olfactory repertoire, an example of “reductive evolution”, as its sensory task is restricted mainly to locating the host’s nest (Hansson and Stensmyr 2011). Similarly, contractions in cytochrome P450 (CYP) genes, such as OG0000009, OG0008679, and OG0000025, reflect the reduced selective pressure for detoxification, since adults are non-foraging and largely solitary, reducing direct exposure to xenobiotics, though the extent to which larval food processing by the host may contribute to reduced detoxification demands remains to be investigated (Feyereisen 1999; Alves-dos-Santos et al. 2002; Johnson et al. 2012). Broad reductions in housekeeping genes involved in ATP-binding, protein-binding, and ion transport further suggest that the cleptoparasitic lifestyle, which spares energy by eliminating nest construction, food processing, and extensive foraging, is mirrored at the genomic level through streamlining of high-energy metabolic pathways (Ricardo et al., 2024). Collectively, these patterns illustrate a dynamic balance in *C. waltheriae* between maintaining essential biological functions and reducing genes rendered superfluous by a parasitic mode of life.

### Ecological interactions inferred from associated genomic sequences

The analysis of sequences that did not belong to the *C. waltheriae* genome revealed an association with mites, fungi, and bacteria, reflecting complex ecological interactions. The identification of *Roubikia* sp., a bee-specific mite also associated with the host *Tetrapedia diversipes, is particularly noteworthy*. This suggests a potential horizontal transmission between host and parasite, consistent with the ecology of Roubikia mites (Cordeiro et al. 2011), which potentially facilitates mite dispersal across host nests. Additionally, the detection of fungi such as *Cokeromyces recurvatus, Malassezia globosa*, and *Trichosporon inkin* indicates significant environmental exposure, as these fungi are commonly found in human-associated and plant-associated habitats. The presence of bacteria including *Bifidobacterium* and *Lactobacillus*, typical members of bee gut microbiomes, suggests a conserved microbiological profile among bees. These findings underscore the importance of incorporating non-target sequence analyses not only to assess genome assembly quality but also to gain insight into the ecological networks and species interactions surrounding specialized taxa such as *C. waltheriae* (Cordeiro et al. 2011; Findley et al. 2013; Roque et al. 2024).

## Conclusions

The genome of *Coelioxoides waltheriae* provides a key reference for understanding the genetic basis of cleptoparasitism in bees. Its compact size results primarily from a reduction in repetitive elements, while core gene content remains comparable to other bees, preserving essential developmental and regulatory functions. Gene family analyses reveal expansions in cuticle and transposable element-related genes, likely facilitating host infiltration and adaptation, alongside contractions in sensory, metabolic, and detoxification genes, reflecting reduced ecological and energetic demands of a parasitic lifestyle. Additionally, non-target sequences analyses highlight interactions with mites, fungi, and bacteria, providing insights into the species’ ecological context. Altogether, this high-quality assembly expands the available genomic resources for solitary bees and offers new opportunities for comparative studies on the evolution of parasitism in Hymenoptera.

## Methods

### DNA extraction, long-read sequencing and processing

Adult cleptoparasitic females of *Coelioxoides waltheriae* were collected at the end of November 2021, a period when the species is actively nesting, on the campus of the University of São Paulo (23° 33′ 53.2″ S 46° 43′ 51.7″ W). Specimens were captured using collecting tubes while flying and searching for nests of *Tetrapedia diversipes* to lay their eggs. They were then placed at -20ºC for euthanasia and stored in liquid nitrogen for later analysis. The whole body of a single individual was used for DNA extraction using the Promega Wizard® HMW DNA Extraction Kit, following the manufacturer’s instructions with two modifications: we used both ethanol and isopropanol at -20ºC, and samples were incubated at -20ºC during DNA precipitation instead of at room temperature. The DNA samples were quantified through a spectrophotometer (NanoDrop× Lite - Thermo Fisher Scientific, Wilmington, Delaware, USA) and shipped to Macrogen (South Korea) for library preparation and sequencing.

The library was constructed using the protocol of SMRTbell™ Template Preparation, containing inserts from ∼250 to 20,000+ bp. The sequencing was performed on a Pacific Biosciences (PacBio) Sequel II platform and its quality was assessed with LongQC v.1.2(Fukasawa et al. 2020).

### Nuclear Genome Assembly

The reads were used for the assembly process of the nuclear genome. Prior to genome assembly, k-mer counting (k = 21) was performed using Jellyfish v2.3.1 (Marçais and Kingsford 2011), and genome characteristics such as expected genome size, heterozygosity and repetitive content were inferred with GenomeScope2 (Vurture et al. 2017). Assembly was performed using Flye v2.9.5-b1801 (Kolmogorov et al. 2019) pipeline, set up for PacBio HiFi reads and scaffold mode. The quality of the final assembly was assesed using Quast v2.5.2 (Gurevich et al. 2013), BloobToolKit v4.4.5 (Challis et al. 2020) and BUSCO v5.8 (Tegenfeldt et al. 2025) tools for the assembly metrics, contamination and completeness analysis.

To avoid noise due to diploidy in female bees, we performed haplotype purging using purge_dups 1.2.5 (Guan et al. 2020). The assembly went on another round of polishing for base correction using arrow (incorporated in Pacbio tools gcpp v1.9.0 – Chin et al. 2016). Reads were then aligned to the assembly using pbmm2 and bwa v0.7.17 (Li 2013), and alignment quality was evaluated with Qualimap v2.2.1 (Okonechnikov et al. 2016). All these processes did not considerably affect the BUSCO and QUAST scores. For the final evaluation and visual representation of the assembly, we used BlobToolKit again to identify potential contamination.

### Comparative Genomic Dataset

A total of 149 reference genomes from Anthophila were retrieved from the National Center for Biotechnology Information (NCBI) database, along with the genome of *C. waltheriae*. This dataset was subsequently used for downstream comparative analyses, focusing on genome size variation and phylogenetic relationships among species (Supplementary Table S1).

### Repetitive Elements Analysis and Gene Prediction and Functional Annotation

Repetitive elements for *C. waltheriae* were identified and analyzed using an adapted version of a previously published pipeline (https://github.com/nat2bee/repetitive_elements_pipeline). A comprehensive repeat library was constructed by combining predictions generated independently with RepeatModeler v2.0.5 (Flynn et al. 2019), TransposonPSI (Haas 2010), and LTRharvest v1.6.7 (Ellinghaus et al. 2008). The resulting libraries were merged and filtered for redundancy using USEARCH v11.0.667 (Zhou et al. 2024) to obtain a non-redundant consensus library. Repeat families were subsequently classified with RepeatClassifier v2.0.5 (Flynn et al. 2019), and used for genome soft-masking with RepeatMasker v4.1.7 (Smit et al. 2013). Final summary statistics were generated and processed using a custom script for graphical representation.

The masked genome was used for protein-coding gene prediction with BRAKER3 v.3.0.8, incorporating both GeneMark-ETP and Augustus predictors (Gabriel et al. 2023). RNA-Seq data from *C. waltheriae* (Ricardo et al., 2024) were aligned to the masked assembly as evidence with hisat2 v2.2.1 (Kim et al. 2015), using the “-*dta*” flag as recommended by BRAKER3’s pipeline. Along with the RNA-Seq data, protein sequences from the OrthoDB 11 database (Arthropoda) (Kuznetsov et al. 2023) were used as extrinsic evidence to support gene prediction. The resulting FASTA and GFF files were then used for functional annotation of the predicted protein sequences. First, an eggNOG-mapper v5.2.0 analysis was performed to identify protein domains, gene ontology terms, and biological pathways (MetaCyc and Reactome) (Cantalapiedra et al. 2021). Additionally, unannotated sequences were subsequently searched against the UniProtKB/Swiss-Prot (Bateman et al. 2023) database using Diamond/BlastX v2.1.10.164 (Buchfink et al., 2015). Remaining unannotated sequences were then searched against the UniProtKB/TrEMBL databaseusing Diamond/BlastX.

To identify genes having repetitive elements in their regulatory region and assess their potential effects, we used a customized pipeline. Briefly, the final annotation file (GTF) provides the start and end positions of the predicted genes. Gene coordinates were first retrieved from the genome annotation. Regulatory regions were defined as 3,000 bp upstream and downstream of each gene. Repetitive elements were identified based on the final RepeatMasker GFF3 output, and their positions were intersected with the defined regulatory regions to determine repeat content within these intervals.

### Non-target DNA sequences identification

The PacBio reads were screened for Acari, bacteria, fungi, human, plants, and viruses sequences to identify potential contaminants. Raw reads were aligned to the masked genome assembly using Minimap2 v2.28-r1209 (Li 2018). Unaligned reads were subsequently isolated and aligned against the NCBI non-redundant nucleotide database (downloaded November 17, 2023) using BLASTn v2.14.1+ (Altschul et al. 1990), with an e-value cutoff of 1e-5. For reads withmultiple hits , only the match with the highest bitscore was retained for taxonomic assignment using the TaxontoolKit v0.20.0 (Shen and Ren 2021). For sequences assigned to Acari, bacteria, viruses, and fungi, information on host associations and environmental occurrence was retrieved from the NCBI database; when unavailable, relevant information was obtained from the literature. Taxa reported to be associated with bees were classified as potentially bee-associated, whereas those linked to other hosts or environmental sources (including human-associated taxa) were considered likely non–bee-associated. Sequences assigned to plant taxa were listed as possible sources of floral resources.

### Orthology Inference

Protein sequence data were obtained from NCBI/GenBank and curated to retain only species with BUSCO single-copy completeness ≥ 90%. A data set composed of *C. waltheria*e along with 38 other bee species, representing six of the seven described bee families (Apidae, Megachilidae, Halictidae, Colletidae, Andrenidae and Melitidae) were built; three wasps (*Ampulex compressa, Cerceris rybyensis* and *Vespula vulgaris* - the first two within Apoidea superfamily and the last from Vespidae), were included as external groups (Supplementary Table S2). To avoid annotation-derived isoform redundancy, sequences were clustered using CD-HIT v4.8.1 (Fu et al. 2012) at 70% similarity, minimizing biases in orthogroup inference and expansion/contraction analysis (see below). Ortholog identification and clustering were performed using OrthoFinder v.2.5.5 (Emms and Kelly 2015) with default parameters. BUSCO v6, in combination with the BUSCO-Phylogenomics (McGowan 2026) pipeline, was used to identify single-copy orthologs across species and construct a supermatrix with 90% occupancy. The sequences within each orthogroup were aligned using MUSCLE v3.8.1551 (Edgar 2004), and poorly aligned regions were trimmed with trimAL v1.4.1 (Capella-Gutiérrez et al. 2009). A maximum likelihood phylogeny was then inferred with IQ-TREE v2.1.3 (Minh et al. 2020) using 1,000 ultrafast bootstrap replicates. To minimize potential biases in model assumptions, we applied the *--symtest-remove-bad* option to exclude loci violating the stationarity, reversibility, and homogeneity assumptions. The resulting phylogram was transformed into a time-calibrated ultrametric tree with calibration points obtained from TimeTree5 (Kumar et al. 2022), setting the root in 179 Mya. The final tree was visually inspected and annotated using iTOL v7 (Letunic and Bork 2024).

### Gene families size changes

We analyzed gene family expansions and contractions across the phylogenetic tree using CAFE v5 (Mendes et al. 2021). To reduce potential biases, the largest gene families identified from the OrthoFinder output were removed using the CAFE script *clade_and_size_filter*.*py*. Next, we estimated key parameters for running CAFE. First, we inferred an error parameter using the original OrthoFinder gene family counts to account for possible genome assembly inaccuracies. This was done by running CAFE with the *-e* flag, which estimates a global error model from the dataset. Next, different k values (from 1 to 10) were tested to identify the model with the highest likelihood. Finally, multiple replicates were run using the selected k value (k = 2) to obtain the median lambda estimates, ensuring robust estimates of gene family evolutionary rate. The final tree containing the topology and the rates for orthologs’ evolutionary changes was obtained using Rstudio (Allaire 2012).

After the CAFE5 analysis, the sets of contracted and expanded orthogroups in *C. waltheriae* were used to retrieve representative gene sequences. Because genes within the same orthogroup are expected to share molecular aspects, and may retain similar or related biological functions, a single representative sequence was selected for each orthogroup following a defined priority hierarchy. (1) Sequences from *C. waltheriae*: whenever available, the first sequence listed for each orthogroup was selected. (2) For contracted orthogroups in which gene loss resulted in the complete absence of *C. waltheriae* sequences, the corresponding *Apis mellifera* sequence was used as reference, given its status as a model species for bees. (3) Arbitrary sequence: when neither *C. waltheriae* nor *A. mellifera* had sequences available for a given orthogroup, the first bee sequence available in the orthogroup, regardless of species, was selected.

Each representative sequence was queried against the NCBI non-redundant (nr) protein database using BLASTP (Altschul et al., 1990) to identify the most likely homologs. To ensure robust functional assignments, only hits with the highest bit score and lowest e-value were reatined, and a conservative score threshold (>100) was applied. Orthogroups not following this criterion were classified as undefined or uncharacterized. The Gene Ontology (GO) annotation obtained from eggNOG-mapper enabled the identification of GO terms enriched in the sets of expanded and contracted orthogroups of *C. waltheriae*. GO enrichment analyses were performed separately for expanded and contracted orthogroups using all annotated orthogroups as background. Analyses were conducted in RStudio with the packages topGO (Alexa and Rahnenführer 2025) and clusterProfiler (Yu et al. 2012), and statistical significance was assessed after Benjamini–Hochberg FDR correction (FDR ≤ 0.05).

## Data Acces

All data generated for this study are deposited at NCBI under the BioProject PRJNA955762. Samples used in this study are authorized under the SISBIO project No 81843-1, in collaboration with the Universidade de São Paulo – Brazil, and were exported under the IBAMA permit 23BR045417/DF, respecting the Nagoya agreement.

## Acknowledgments

We thank Suzy Coelho for her technical contribution to the data production.

## Declarations

### Ethics approval and consent to participate

Not applicable.

### Authors contributions

All authors contributed to the manuscript preparation and revision, below are listed other specific contributions. FCD : study design, data production and analysis, manuscript preparation. HM: data production and analysis and manuscript preparation. PCR: data analysis and manuscript preparation. BTM: data analysis. NSA: study design. MCA: study design, funding, manuscript preparation and sutdy coordination.

