## Supplemental Figure 1 for "The high-quality genome assembly of Coelioxoides waltheriae (Apidae: Nomadinae) reveals gene family dynamics and evolutionary shifts related to its cleptoparasitic lifestyle"

**Fig. S1** GenomeScope profile plots of absolute (top graph) and log-transformed (bottom graph) k-mer frequency distributions, at a k-mer length of 21, built based on the PacBio HiFi long-reads.

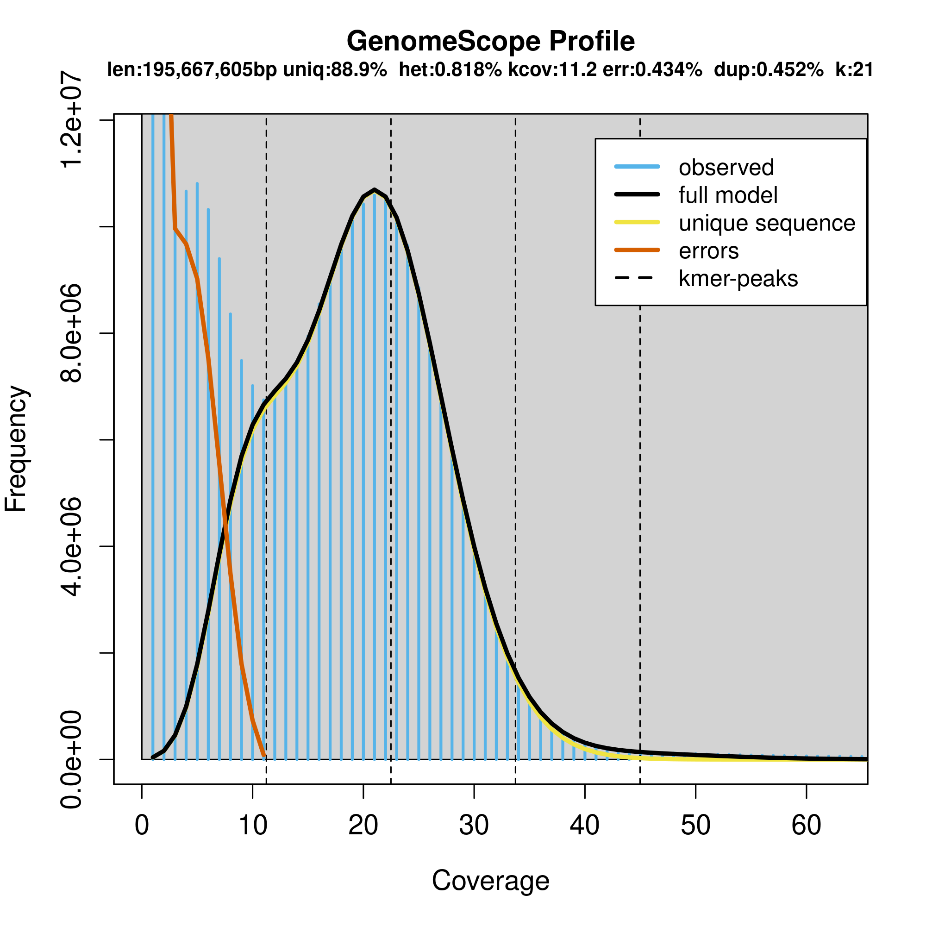

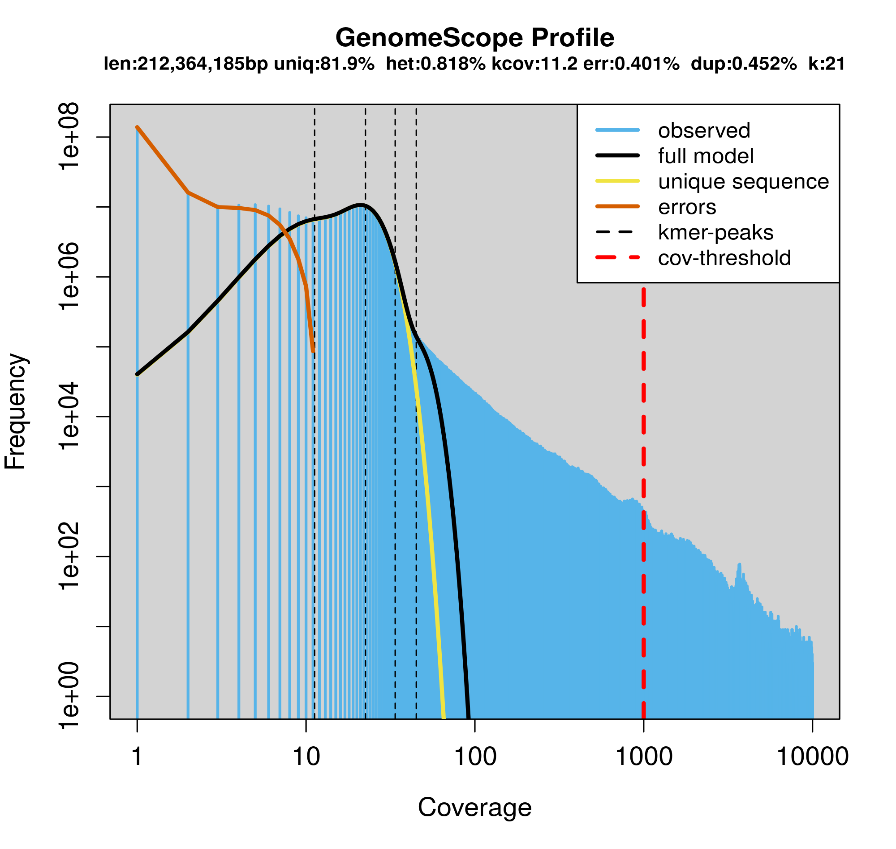
