## Supplemental Table 3 for "The high-quality genome assembly of Coelioxoides waltheriae (Apidae: Nomadinae) reveals gene family dynamics and evolutionary shifts related to its cleptoparasitic lifestyle"

| **Supplementary Table S3** | | | |
| --- | --- | --- | --- |
|  | **Number of elements** | **Length occupied** | **Percentagem of sequence** |
| Retroelements | 5346 | 1280203 bp | 0.66 % |
| SINEs | 0 | 0 | 0.00 % |
| Penelopes | 0 | 0 | 0.00 % |
| LINEs | 1077 | 319329 bp | 0.16 % |
| CRE/SLACS | 0 | 0 | 0.00 % |
| L2/CR1/Rex | 428 | 87739 bp | 0.05 % |
| R1/LOA/Jockey | 486 | 128483 bp | 0.07 % |
| R2/R4/NeSL | 73 | 73191 bp | 0.04 % |
| RTE/Bov-B | 5 | 2179 bp | 0.00 % |
| L1/CIN4 | 0 | 0 | 0.00 % |
| LTR elements | 4269 | 960874 bp | 0.49 % |
| BEL/Pao | 27 | 29123 bp | 0.01 % |
| Ty1/Copia | 560 | 149848 bp | 0.08 % |
| Gypsy/DIRS1 | 1926 | 577449 bp | 0.30 % |
| Retroviral | 1685 | 192508 bp | 0.10 % |
| DNA transposons | 43370 | 5719986 bp | 2.94 % |
| hobo-Activator | 198 | 45988 bp | 0.02 % |
| Tc1-IS630-Pogo | 9380 | 1635822 bp | 0.84 % |
| En-Spm | 0 | 0 | 0.00 % |
| MULE-MuDR | 0 | 0 | 0.00 % |
| PiggyBac | 13493 | 1805337 bp | 0.93 % |
| Tourist/Harbinger | 106 | 11144 bp | 0.01 % |
| Other (Mirage, P- element, Transib) | 0 | 0 | 0.00 % |
| Rolling-circles | 1429 | 279214 bp | 0.14 % |
| Unclassified: | 103627 | 16572088 bp | 8.51 % |
| Total interspersed repeats: | | 23572277 bp | 12.10 % |
| Small RNA: | 221 | 62275 bp | 0.03 % |
| Satellites: | 533 | 62390 bp | 0.03 % |
| Simple repeats: | 65988 | 3662706 bp | 1.88 % |
| Low complexity: | 15321 | 897059 bp | 0.46 |
