## Supplemental Table 7 for "The high-quality genome assembly of Coelioxoides waltheriae (Apidae: Nomadinae) reveals gene family dynamics and evolutionary shifts related to its cleptoparasitic lifestyle"

| **Table S9.** Overview of the predicted orthogroup changes, expansions (2^nd^ to 4^th^ columns) and contractions (5^th^ to 7^th^ columns), for the genome of each insect species analyzed. Numbers between parenthesis in the 2^nd^ and 5^th^ columns indicate the number of orthogroups found to be rapidly expanding and contracting, respectively. Positive or negative numbers in the 9^th^ column indicate an overall expansion or contraction, respectively. | | | | | | | | |
| --- | --- | --- | --- | --- | --- | --- | --- | --- |
| **Species** | **Expanded OGs** | **Orthologs gained** | **Orthologs**  **/expansion** | **Contracted OGs** | **Orthologs**  **lost** | **Orthologs/**  **contraction** | **No OGs**  **change** | **Avg.**  **expansion** |
| *Ampulex compressa<2>* | 379 (84) | 445 | 1,17 | 198 (115) | 209 | 1,05 | 8614 | 0,025677293 |
| *Andrena bucephala<6>* | 444 (136) | 556 | 1,25 | 691 (254) | 711 | 1,02 | 8056 | -0,016864324 |
| *Andrena dorsata<5>* | 964 (110) | 1055 | 1,09 | 1267 (476) | 1298 | 1,02 | 6960 | -0,026438908 |
| *Anthophora plagiata<23>* | 669 (121) | 739 | 1,10 | 965 (403) | 984 | 1,01 | 7557 | -0,026656512 |
| *Anthophora quadrimaculata<24>* | 703 (115) | 772 | 1,09 | 351 (130) | 353 | 1,00 | 8137 | 0,045588075 |
| *Apis cerana<31>* | 696 (144) | 819 | 1,17 | 229 (81) | 231 | 1,00 | 8266 | 0,063975628 |
| *Apis mellifera<32>* | 829 (92) | 930 | 1,12 | 341 (178) | 347 | 1,01 | 8021 | 0,063431618 |
| *Bombus pratorum<34>* | 159 (47) | 174 | 1,09 | 354 (222) | 361 | 1,01 | 8678 | -0,020345991 |
| *Bombus sylvestris<36>* | 631 (176) | 749 | 1,18 | 484 (130) | 486 | 1,00 | 8076 | 0,028614949 |
| *Bombus terrestris<33>* | 203 (62) | 263 | 1,29 | 610 (188) | 635 | 1,04 | 8378 | -0,040474377 |
| *Bombus vestalis<35>* | 659 (127) | 750 | 1,13 | 266 (95) | 271 | 1,01 | 8266 | 0,052116201 |
| *Ceratina calcatara<27>* | 764 (122) | 881 | 1,15 | 478 (176) | 494 | 1,03 | 7949 | 0,042106408 |
| *Cerceris rybyensis<3>* | 730 (129) | 814 | 1,11 | 789 (350) | 807 | 1,02 | 7672 | 0,000761615 |
| *Coelioxoides waltheriae<18>* | 191 (84) | 223 | 1,16 | 259 (145) | 264 | 1,01 | 8741 | -0,004460886 |
| *Coelioxys conoideus<16>* | 380 (122) | 446 | 1,17 | 254 (111) | 254 | 1 | 8557 | 0,020890001 |
| *Colletes collaris<9>* | 150 (52) | 177 | 1,18 | 641 (255) | 669 | 1,04 | 8400 | -0,053530628 |
| *Colletes gigas<8>* | 246 (77) | 294 | 1,19 | 455 (157) | 464 | 1,01 | 8490 | -0,018496355 |
| *Eufrisea auriceps<30>* | 112 (34) | 143 | 1,27 | 568 (264) | 590 | 1,03 | 8511 | -0,048634534 |
| *Eufrisea mexicana<29>* | 348 (130) | 439 | 1,26 | 586 (177) | 604 | 1,03 | 8257 | -0,017952345 |
| *Exaerette dentata<28>* | 616 (131) | 658 | 1,06 | 173 (71) | 174 | 1,00 | 8402 | 0,052660211 |
| *Friesiomelitta varia<41>* | 607 (114) | 641 | 1,05 | 159 (71) | 163 | 1,02 | 8425 | 0,052007399 |
| *Habropoda laboriosa<22>* | 105 (42) | 116 | 1,10 | 373 (189) | 378 | 1,01 | 8713 | -0,028506147 |
| *Holcopasites calliopsidis<19>* | 219 (50) | 265 | 1,21 | 269 (88) | 272 | 1,01 | 8703 | -0,000761615 |
| *Hylaeys volcanicus<7>* | 190 (39) | 219 | 1,15 | 852 (169) | 864 | 1,01 | 8149 | -0,070177347 |
| *Lasioglossum calceatum<11>* | 483 (175) | 643 | 1,33 | 757 (241) | 774 | 1,02 | 7951 | -0,014253074 |
| *Lestrimelitta limao<40>* | 360 (112) | 444 | 1,23 | 157 (87) | 163 | 1,03 | 8674 | 0,030573387 |
| *Macropis europaea<4>* | 381 (129) | 469 | 1,23 | 411 (187) | 422 | 1,02 | 8399 | 0,005113698 |
| *Megachile rotundata<15>* | 170 (46) | 177 | 1,04 | 566 (211) | 586 | 1,03 | 8455 | -0,044500054 |
| *Megalopta genalis<10>* | 218 (122) | 273 | 1,25 | 281 (125) | 282 | 1,00 | 8692 | -0,000979219 |
| *Melipona quadrifasciata<39>* | 121 (42) | 140 | 1,15 | 524 (244) | 541 | 1,03 | 8546 | -0,043629638 |
| *Melipona scutelaris<38>* | 228 (87) | 265 | 1,16 | 136 (68) | 137 | 1,00 | 8827 | 0,013926667 |
| *Nomada flavoguttata<20>* | 171 (68) | 182 | 1,06 | 220 (109) | 223 | 1,01 | 8800 | -0,004460886 |
| *Nomada lathburiana<21>* | 582 (130) | 643 | 1,10 | 263 (91) | 271 | 1,03 | 8346 | 0,040474377 |
| *Osmia bicornis<14>* | 596 (126) | 657 | 1,10 | 278 (100) | 283 | 1,01 | 8317 | 0,040691981 |
| *Protosiris gigas<17>* | 529 (100) | 575 | 1,08 | 369 (158) | 380 | 1,02 | 8293 | 0,021216407 |
| *Sphecodes monilicornis<12>* | 593 (129) | 679 | 1,14 | 220 (85) | 228 | 1,03 | 8378 | 0,049069742 |
| *Stelis phaeoptera<13>* | 77 (32) | 94 | 1,22 | 501 (273) | 506 | 1,00 | 8613 | -0,044826461 |
| *Tetragonisca angustula<42>* | 214 (59) | 226 | 1,05 | 219 (89) | 227 | 1,03 | 8758 | -0,000108802 |
| *Tetragonula carbonaria<37>* | 178 (69) | 192 | 1,07 | 182 (80) | 191 | 1,04 | 8831 | 0,000108802 |
| *Tetrapedia diversipes<25>* | 159 (59) | 186 | 1,16 | 472 (257) | 497 | 1,05 | 8560 | -0,03383745 |
| *Vespula vulgaris<1>* | 490 (201) | 560 | 1,14 | 200 (96) | 206 | 1,03 | 8501 | 0,03851594 |
| *Xylocopa violacea<26>* | 138 (56) | 340 | 2,46 | 385 (187) | 390 | 1,01 | 8668 | -0,005440104 |

| Taxon | Association |
| --- | --- |
| Acari  *Dermatophagoides sp.*  *Sarcoptes scabiei*  *Roubikia sp.* | human human  *C. waltheriae* and *Tetrapedia diversipes* |
| Fungi  *Fulvia fulva*  *Botrytis cinerea,*  *Lasiodiplodia theobromae,*  *Aspergillus nomiae*  *Cokeromyces recurvatus*  *Malassezia globos*a  *Trichosporon inkin* | plant pathogen plant pathogen plant pathogen plant pathogen  human  human  human |
| Bacteria  *Bifidobacterium sp.*  *Bombella sp.*  *Lactobacillus sp.*  *Melissococcus plutonius*  *Parasaccharibacter apium*  *Parasaccharibacter sp.*  *Snodgrassella alvi*  *Snodgrassella communis*  *Aristophania vespae*  *Oecophyllibacter saccharovorans*  *Proteus vulgaris*  *Other* | bee-host associated  bee-host associated  bee-host associated  bee-host associated  bee-host associated  bee-host associated  bee-host associated  bee-host associated    insect-associated (non Apidae)    human, plant and environmental bacteria |
| Human |  |
| Hymnoptera  Nomadinae |  |
| Virus |  |
