## Supplementary figures and images for "The high-quality genome assembly of Coelioxoides waltheriae (Apidae: Nomadinae) reveals gene family dynamics and evolutionary shifts related to its cleptoparasitic lifestyle"

### Supplemental Figure 2

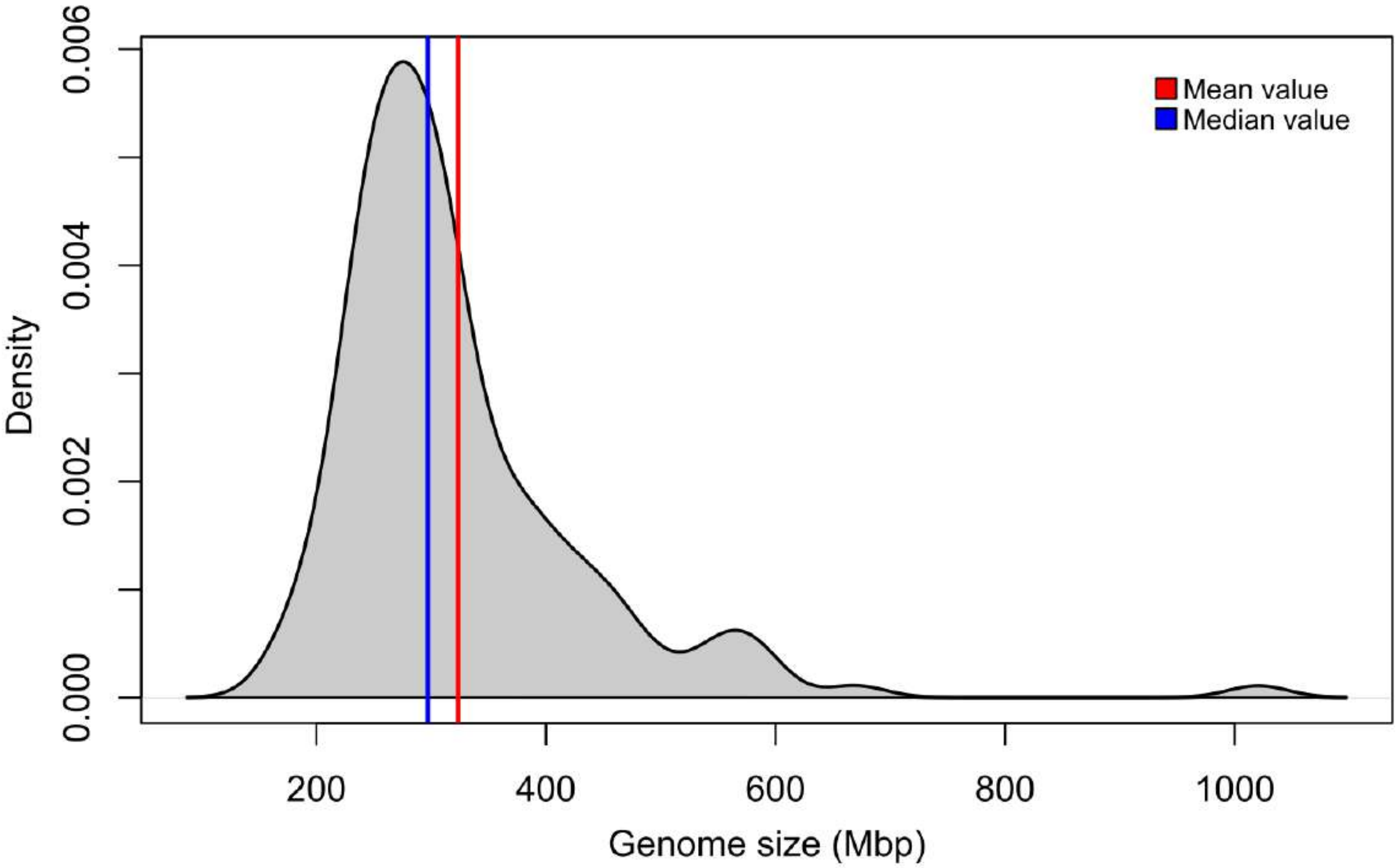
